# Competition between microtubule-associated proteins directs motor transport

**DOI:** 10.1101/180935

**Authors:** Brigette Y. Monroy, Danielle L. Sawyer, Bryce E. Ackermann, Melissa M. Borden, Tracy C. Tan, Kassandra M. Ori-McKenney

## Abstract

Within cells, numerous motor and non-motor microtubule-associated proteins (MAPs) simultaneously converge on the microtubule lattice. How the binding activities of non-motor MAPs are coordinated and how they contribute to the balance and distribution of microtubule motor transport is unknown. Here, we examine the relationship between MAP7 and tau due to their antagonistic effects on neuronal branch formation and kinesin motility in vivo^1–8^. We find that MAP7 and tau compete for binding to microtubules, and determine a mechanism by which MAP7 displaces tau from the lattice. In striking contrast to the inhibitory effect of tau, MAP7 promotes kinesin-based transport in vivo and strongly enhances kinesin-1 binding to the microtubule in vitro, providing evidence for direct enhancement of motor motility by a MAP. In contrast, both MAP7 and tau strongly inhibit kinesin-3 and have no effect on cytoplasmic dynein, demonstrating that MAPs exhibit differential control over distinct classes of motors. Overall, these results reveal a general principle for how MAP competition dictates access to the microtubule to determine the correct distribution and balance of molecular motor activity.

---

The balance of intracellular transport is essential for the structural and functional organization of a cell. The microtubule motors, kinesin-1 and cytoplasmic dynein, drive cellular cargoes towards microtubule plus-ends and minus ends, respectively. Mutations in either motor pathway disrupt the balance of transport and cause a range of diseases^9,10^. Motors must navigate a crowded microtubule lattice that is decorated with non-motor MAPs. These MAPs play various roles in regulating microtubule dynamics, turnover, and stability, as well as in influencing molecular motor transport^3,11–13^. The Alzheimer’s disease related MAP, tau (MAPT), is highly enriched in neuronal axons and inhibits kinesin-1 motility in vivo and in vitro, but has less of an effect on dynein-based movement^3–8^. How plus-end directed transport is accomplished in the tau-rich axonal environment is therefore an outstanding question, though the effects of tau on other classes of kinesin motors have not been tested. One MAP that appears to be antagonistic to tau is MAP7 (EMAP-115, ensconsin)^14,15^. The knockdown of MAP7 or tau in neurons produces opposite axonal branching phenotypes^1,2^. In addition, MAP7 is important for kinesin-based cargo transport and nuclear positioning in vivo^16–18^; however, the molecular mechanism underlying this relationship is unclear. How tau and MAP7 activities are coordinated on the surface of individual microtubules in cells, and the functional consequence of MAP distributions on microtubule motor motility remain open questions.

To approach this problem, we first examined the localization patterns of MAP7 and tau in mature *Drosophila* peripheral nervous system neurons in vivo and DIV4 primary mouse neuronal cultures. In both systems, we found that MAP7 localized within both dendrites and axons, while tau was predominantly restricted to the axons, as previously reported (Fig. 1a-d)^2,19^. Based on their overlapping spatial patterns within neurons, we wanted to investigate whether MAP7 and tau could bind simultaneously to individual microtubules. Using total internal reflection fluorescence microscopy (TIRF-M), we imaged purified fluorescently labeled human 2N4R tau and full-length MAP7 (Supplemental Fig. 1) binding to taxol-stabilized microtubules. We first mixed equimolar concentrations of both MAPs in the flow chamber and observed that tau is strongly excluded from sites of MAP7 enrichment, and vice versa (Fig. 2a-d). A five-fold excess of MAP7 completely abolished tau binding to the microtubule (Fig. 2a-b). Conversely, even at 100-fold excess of tau, MAP7 remained tenaciously bound to the microtubule in distinct patches that were devoid of tau (Fig. 2c-d). These experiments suggest that MAP7 and tau compete for binding to the microtubule surface.

**Figure 1.**
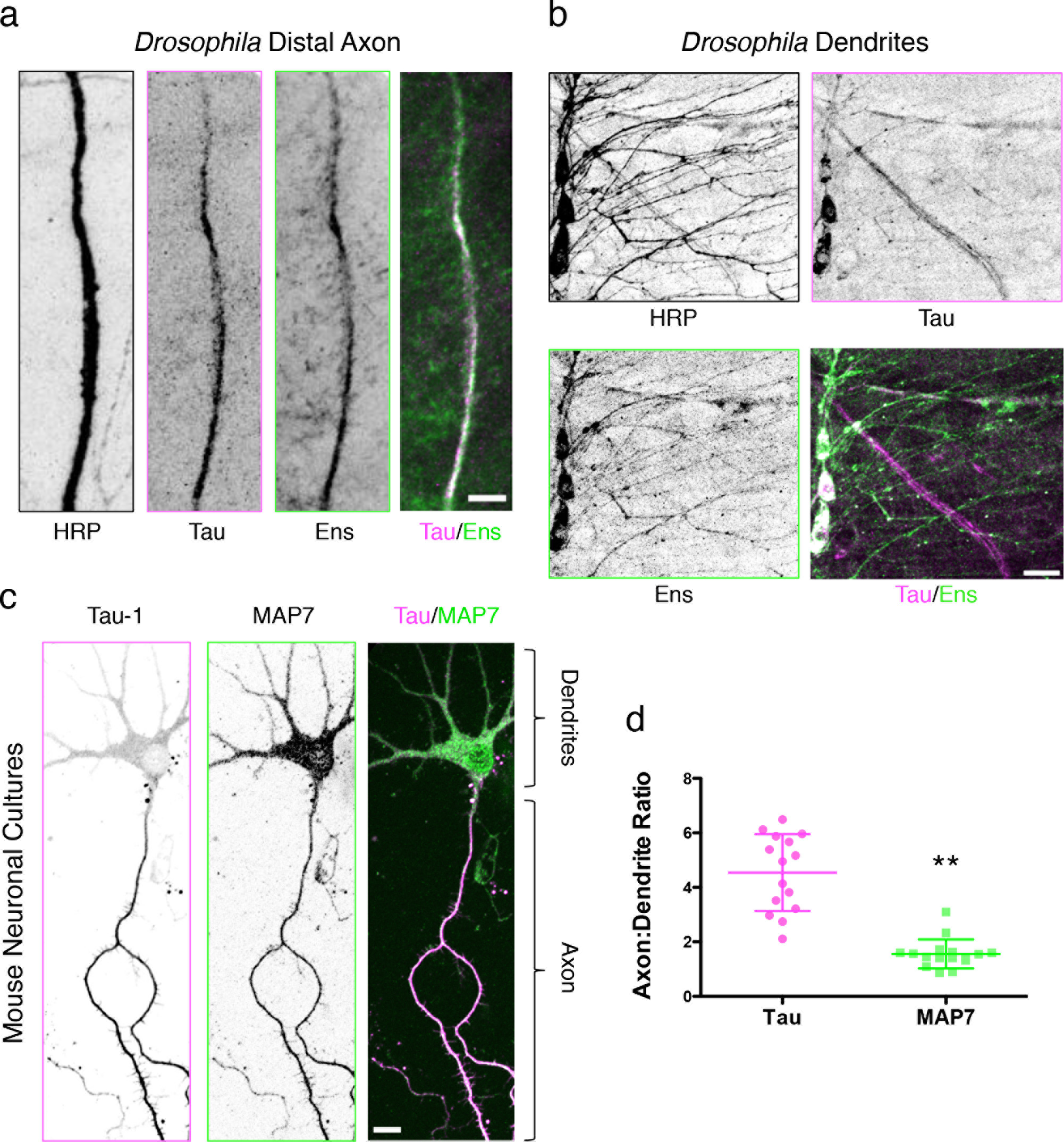
MAP7 and tau exhibit overlapping localization patterns in *Drosophila* and mammalian neurons. Immunohistochemical staining of larval fillets from an endogenous tau-GFP fly line with antibodies against HRP, GFP, and ensconsin reveals that (**a**) tau and ensconsin are expressed within axons, and (**b**) ensconsin, but not tau, is expressed in the dendrites of peripheral nervous system neurons. Scale bars are 10 and 15 μm for (**a**) and (**b**), respectively. (**c**) Immunocytochemistry of mouse DIV4 neuronal cultures with antibodies against dephospho-tau (Tau-1) and MAP7. Scale bars are 20 μm. (**d**) Quantification of the axon to dendrite fluorescence intensity ratio for MAP7 is 1.5 **±** 0.5, indicating a near uniform distribution of MAP7 throughout the neuron, while the ratio for tau is 4.5 **±** 1.4, indicating a five-fold enrichment of tau in the axon (*P* < 0.0001, n = 15 and 16 neurons for tau and MAP7, respectively from two independent cultures).

**Figure 2.**
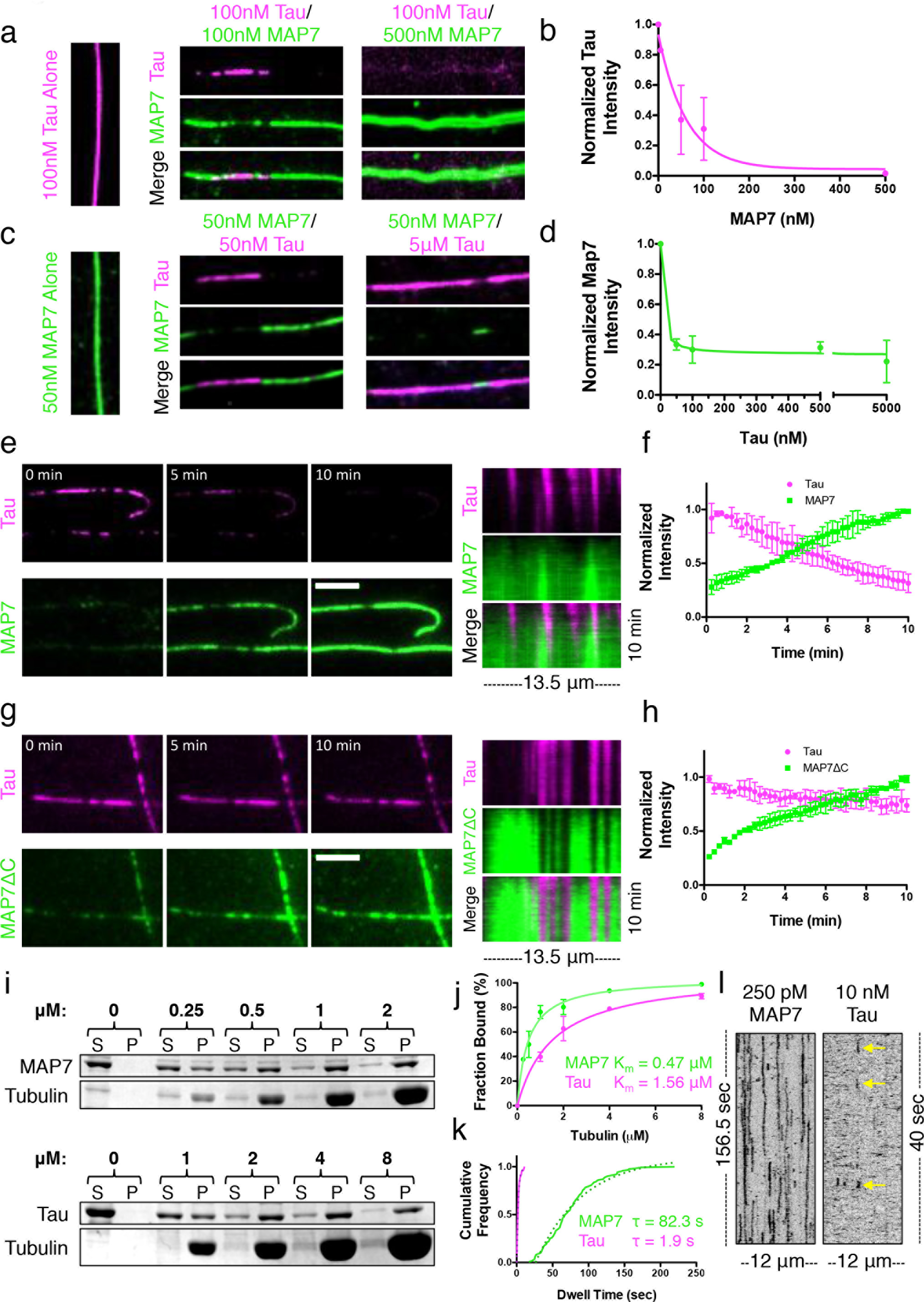
MAP7 and tau compete for binding on the microtubule lattice. (**a**) TIRF-M images of mTagBFP-tau (pink) binding to taxol-stabilized microtubules in the absence and presence of increasing concentrations of sfGFP-MAP7 (green). (**b**) Quantification of tau fluorescence intensity as a function of increasing MAP7 concentration (n = 320 microtubules quantified from two independent trials). (**c**) Images of sfGFP-MAP7 (green) binding to taxol-stabilized microtubules in the absence and presence of increasing concentrations of mTagBFP-tau (pink). (**d**) Quantification of MAP7 fluorescence intensity as a function of increasing tau concentration (n = 228 microtubules quantified from three independent trials). All images in (**a**) and (**c**) are 10.3 μm in length. (**e**) Successive movie frames of 50 nM mTagBFP-tau and 50 nM sfGFP-MAP7 on the microtubule reveals the competition between MAP7 and tau. Scale bar is 5 μm. The corresponding kymograph to the right depicts the invasion and displacement of tau patches by MAP7 over 10 min (images were acquired at 15 s intervals). (**f**) Quantification of tau and MAP7 fluorescence intensity over 10 min (n = 6 microtubules quantified per 15 s interval from two independent trials). (**g**) Successive movie frames of 50 nM mTagBFP-tau and 50 nM sfGFP-MAP7ΔC show that without its acidic C-terminus, MAP7 can no longer evict tau from the microtubule. Scale bar is 5 μm. The corresponding kymograph to the right depicts the behavior of tau and MAP7ΔC over 10 min (images were acquired at 15 s intervals). (**h**) Quantification of tau and MAP7ΔC fluorescence intensity over 10 min (n = 6 microtubules quantified per 15 s interval from two independent trials). (**i**) Coomassie Blue-stained SDS-Polyacrylamide gel electrophoresis (SDS-PAGE) shows the binding behavior of 500 nM MAP7 or 500 nM tau in the presence of increasing concentrations of taxol-stabilized microtubules. (**j**) Results from three separate microtubule co-sedimentation experiments were plotted and fit to a Michaelis-Menten equation producing a KD **±** s.d of 0.47 **±** 0.06 μM for MAP7 and 1.56 **±** 0.16 μM for tau (*P* = 0.0004). (**k**) Graph depicting the cumulative frequency of MAP7 (decay constant, *τ* = 82.3 **±** 11.2 s; n = 206 events from n = 39 microtubules from two independent trials) and tau (decay constant, *τ* = 1.9 **±** 0.1 s; n = 142 events from n = 16 microtubules from two independent trials) dwell times fit to a one phase exponential decay (R^2^ = 0.980 and 0.995 for MAP7 and tau, respectively; *P* < 0.0001). (**l**) Corresponding representative kymographs for 250 pM sfGFP-MAP7 and 10 nM sfGFP-tau dwell times on taxol-stabilized microtubules (frame rates are 2 f/s and 7.8 f/s for MAP7 and tau, respectively due to the differences in binding dynamics). Yellow arrows indicate the short tau dwells. Means **±** s.d. are plotted for all graphs in (**b**), (**d**), (**f**), (**h**), and (**j**). For all experiments, similar results were obtained with sfGFP-tau and TagRFP-MAP7 and repeated with at least two separate protein preparations for each protein.

We next performed time-lapse imaging of MAP7 and tau dynamics on microtubules, and observed that while tau initially bound microtubules more rapidly than MAP7, over longer time periods, MAP7 accumulated and displaced tau from the lattice (Fig. 2e-f). There are two non-mutually exclusive possibilities for how MAP7 and tau could compete for the microtubule: 1) they share overlapping binding sites on the microtubule, or 2) domains of each MAP not involved in microtubule binding could play a role in hindering the binding of the other. We tested these possibilities by purifying a truncated version of MAP7 that contains the microtubule-binding domain, but lacks a conserved negatively charged C-terminal region (-10.7 net charge, MAP7AC). We find that although MAP7AC still accumulates on the microtubule in patches that exclude tau, it is unable to completely evict tau from the microtubule, indicating that MAP7 and tau do share overlapping binding sites, but the negatively charged C-terminal domain of MAP7 plays a secondary role in evicting tau molecules that are already associated with the lattice (Fig. 2g-h).

Next, we evaluated the microtubule binding behavior of MAP7 and tau. Microtubule cosedimentation assays revealed that MAP7 bound microtubules with an apparent K_D_ of 0.46 **±** 0.06 μM, while tau displayed a 3-fold higher K_D_ of 1.56 **±** 0.16 μM in our conditions (Fig. 2i-j). These values are similar to previous in vitro results and suggest that MAP7 has a higher microtubule binding affinity than tau^14,20^. Nonetheless, we observed tau binding to microtubules before MAP7 in our TIRF-M assays, which could be attributed to the cooperativity exhibited by tau^21^. Single molecule dwell time analysis revealed that MAP7 molecules bind the microtubule ~40-times longer than tau molecules in our conditions (82.3 s vs. 1.9 s; Fig. 2k-l), though these binding dynamics are likely modulated by external factors in vivo, such as by phosphorylation^22–24^. These results indicate that MAP7 and tau compete for a common binding site on the microtubule and MAP7 is able to invade tau rich regions of the microtubule through a mechanism that involves its negatively charged C-terminal domain, its higher microtubule binding affinity, and longer lattice residency times.

To determine the functional consequence of the MAP7-tau competition, we analyzed the effects of tau and ensconsin, the *Drosophila* homologue of MAP7, overexpression in larval dendritic arborization (DA) neurons. Tau overexpression led to fewer branches, while ensconsin overexpression caused an increase in the number of dendrite branches and total dendrite length (Fig. 3a-c). Overexpression of both tau and ensconsin still led to an increase in the number of branches and total branch length (Fig. 3a-c), consistent with the ability of MAP7 to outcompete tau in vitro. Interestingly, the increase in proximal branching that results from ensconsin overexpression phenocopies dynein knockdown in these neurons^25^, indicating this phenotype may be the result of an imbalance in motor transport. We examined the localization pattern of Golgi outposts because this cargo is bidirectionally transported in dendrites and is important for branch formation within the DA neurons^26,27^. In control neurons, Golgi outposts are enriched within the cell body, within the primary and secondary branches, at branchpoints, and at distal tips (Fig. 3d)^28^. Overexpression of ensconsin redistributed Golgi outposts from the primary and secondary branches to the cell body and the extreme distal tips of dendrite branches (Fig. 3d-e). Based on the orientation of microtubules within these neurons, these areas are likely enriched in microtubule plus-ends (Fig. 3f)^29^. These data indicate that overexpression of ensconsin possibly influences the balance of transport in favor of kinesin, driving plus-end directed movement of kinesin cargoes such as Golgi outposts. Accumulation of Golgi outposts near the cell body may facilitate nascent branch formation in the proximal region via microtubule nucleation^28,30^, providing an explanation for the phenotype resulting from ensconsin overexpression.

**Figure 3:**
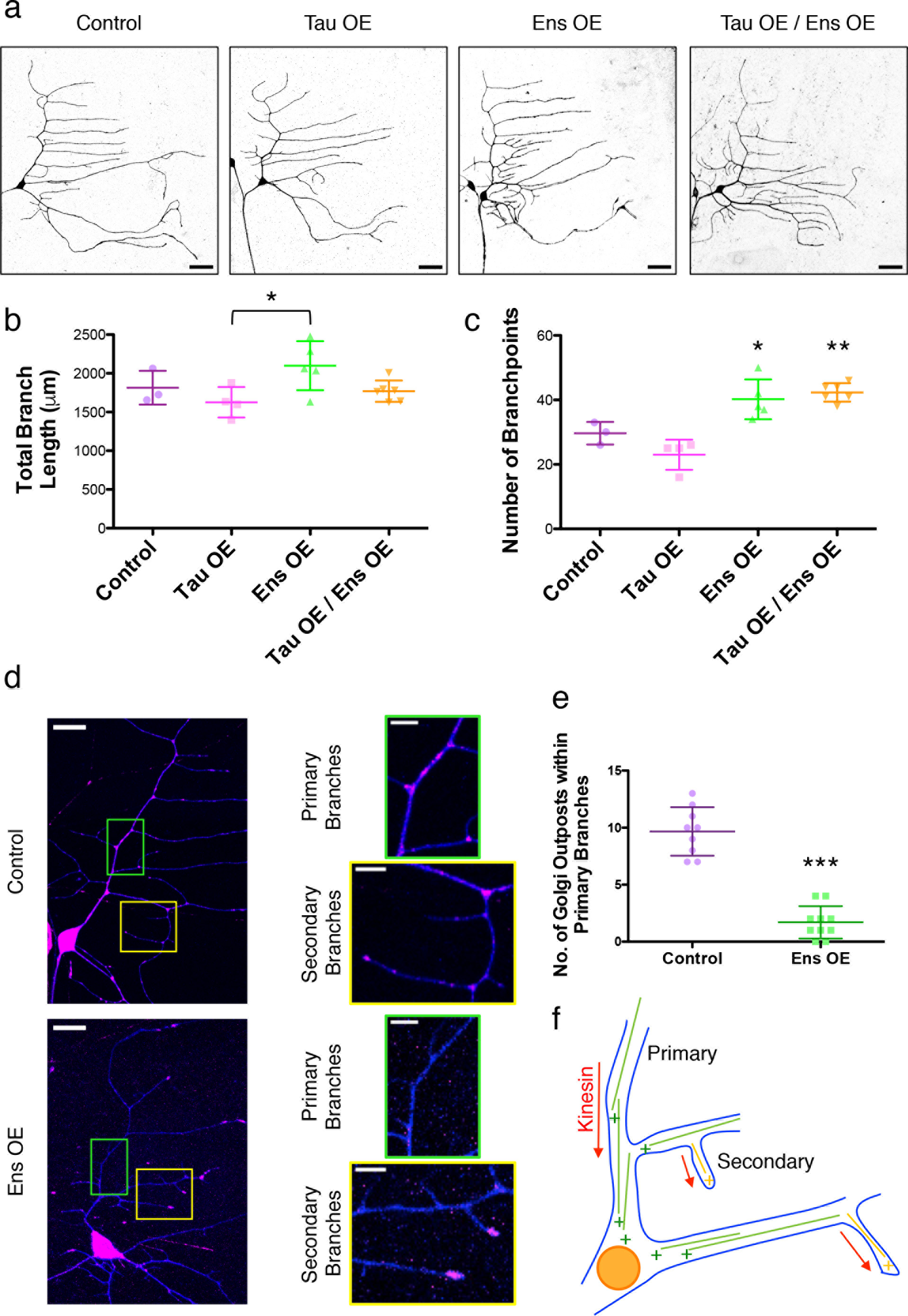
*Drosophila* ensconsin enhances kinesin transport in vivo. (**a**) Dendrite morphologies of control, *221-Gal4>*UAS-*tau* overexpression (OE), *221-Gal4>*UAS-*ens* OE, and *221-Gal4>*UAS-*tau* OE/UAS-*ens* OE class I DA neurons. Neurons were visualized using *UAS-CD4-RFP* expressed by *221-Gal4*. Scale bars are 30 μm. (**b**) Quantification of the total neuronal branch length for each genotype (means **±** s.d. are 1814 **±** 218, 1703 **±** 151, 2099 **±** 317, and 1769 **±** 137 μm for control, tau OE, ens OE, and tau OE/ens OE, respectively; control vs. tau OE: *P* = 0.5; control vs. ens OE: *P* = 0.2; control vs. tau OE/ens OE: *P* = 0.7; tau OE vs. ens OE: *P* = 0.09) (**c**) Quantification of the number of dendritic branchpoints for each genotype (means **±** s.d. are 29.7 **±** 3.5, 25.3 **±** 0.6, 40.2 **±** 6.2, and 42.3 **±** 2.9 branchpoints for control, tau OE, ens OE, and tau OE/ens OE, respectively; control vs. tau OE: *P* = 0.1; control vs. ens OE: *P* = 0.03; control vs. tau OE/ens OE: *P* = 0.0006; tau OE vs. ens OE: *P* = 0.007). n = 3, 3, 5, and 6 neurons for control, tau OE, ens OE, and tau OE/ens OE, respectively for (**b**) and (**c**). (**d**) Golgi outpost (pink) localization throughout the dendritic arbor (blue) of control and *221-Gal4>UAS-ens* OE class I DA neurons, visualized by *221-Gal4>UAS-ManII-eGFP, UAS-CD4-RFP*. Boxes of zoomed regions reveal primary and secondary branches filled with Golgi outposts in control neurons, but devoid of Golgi outposts in ensconsin OE neurons. Golgi outposts are driven towards the cell body or distal dendrite tips. Scale bars are 15 μm for full neurons and 5 μm for zoomed images. (**e**) Quantification of the number of Golgi outposts per 50 μm of primary branch in control vs. ensconsin OE neurons (means **±** s.d. are 9.7 **±** 2.1 vs. 1.7 **±** 1.4 Golgi outposts for control and ensconsin OE, respectively; *P* < 0.0001; n = 10 neurons for each genotype). All graphs in (**b**), (**c**), and (**e**) are scatterplots of all datapoints with the lines indicating the means **±** s.d. (**f**) Schematic of the orientation of microtubules within the class I DA neurons, indicating that sites of Golgi outpost localization in the ensconsin OE neurons are final destinations for microtubule plus-end transport.

Kinesin-1 has been reported to drive the anterograde transport of intracellular cargoes within the dendrites of the DA neurons^31^, indicating the overexpression of ensconsin could indeed enhance kinesin-1 transport. In addition, previous studies have reported that MAP7 is necessary for kinesin-1 activities in vivo^16–18,32^; however, the molecular mechanism by which MAP7 affects kinesin-1, either directly or indirectly, is unknown. To determine if MAP7 directly affects kinesin-1, we analyzed the dynamics of purified MAP7 and K560, a well-characterized truncated version of human kinesin-1 ^33^, on taxol-stabilized microtubules using TIRF-M. We found that the presence of saturating MAP7 on the microtubule (Supplemental Fig. 2a) dramatically enhanced the amount of K560 on the microtubule 165-fold (Fig. 4a), and increased the landing rate of K560 ~15-fold (from 0.09 **±** 0.11 to 1.34 **±** 0.43 motors per μM per min. Fig. 4b). MAP7 also slightly increased the processivity (from 873.8 **±** 43.3 to 984.3 **±** 87.5 nm) and decreased the velocity of K560 (from 434.0 **±** 111.2 to 327.5 **±** 128.5 nm/sec, Fig. 4c-d) in our assays. To test the effects of MAP7 on K560 enzymatic activity, we measured the basal and microtubule-stimulated ATPase activities of K560 in the presence and absence of MAP7. The basal ATPase activity of K560 was unchanged by MAP7; however, MAP7 lowered the K_mMT_ of K560 from 1.46 **±** 0.21 μM to 0.27 **±** 0.04 μM, indicating that MAP7 increases the affinity of K560 for the microtubule without affecting its ATPase turnover rate (Fig. 4e).

**Figure 4:**
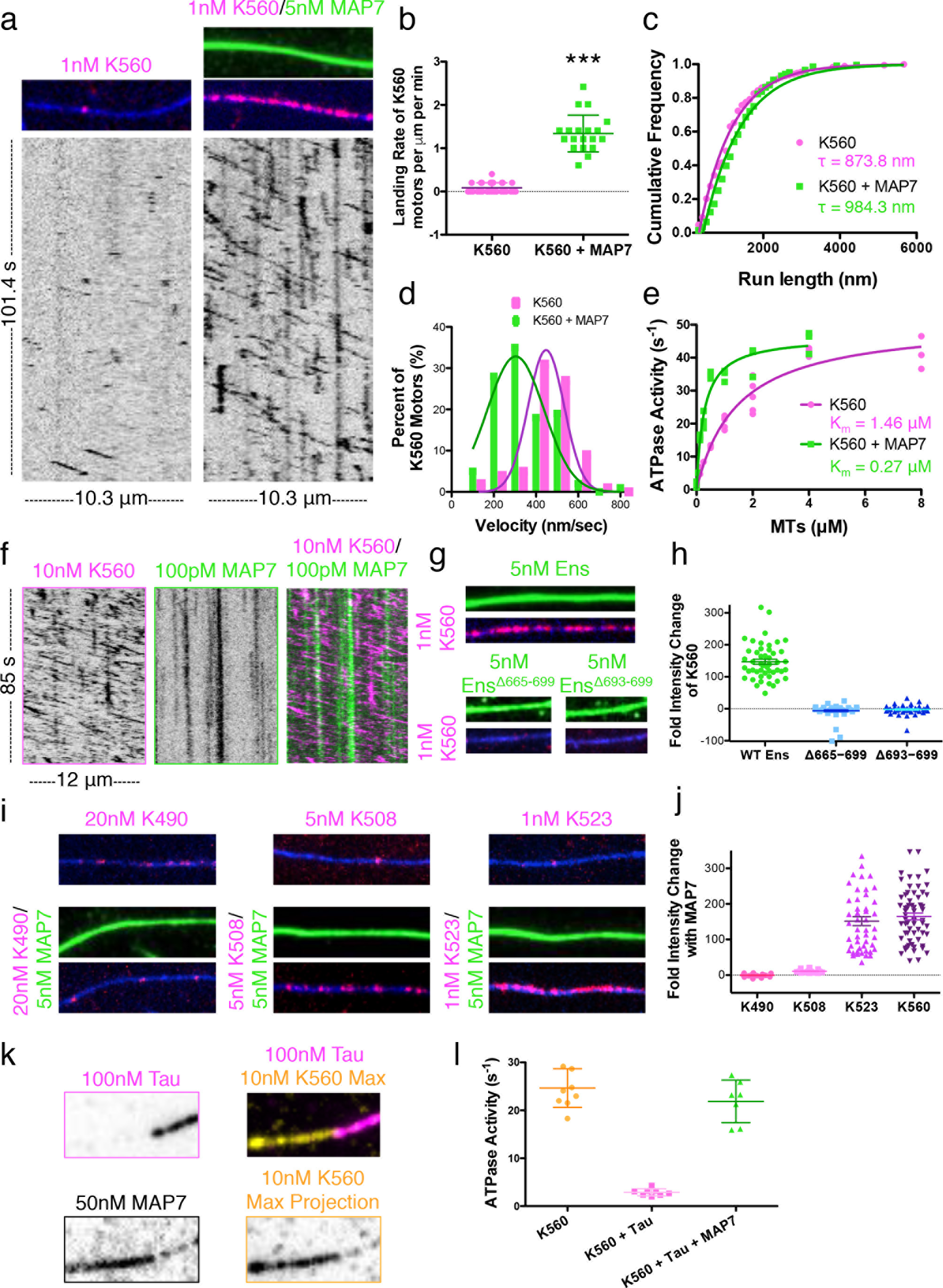
MAP7 directly recruits kinesin-1 to the microtubule. (**a**) TIRF-M images and corresponding kymographs of 1 nM K560-mScarlet (pink) + 1 mM ATP in the absence and presence of 5 nM sfGFP-MAP7 (green). Images are 10.3 μm in length. (**b**) Quantification of the landing rates of 1 nM K560-mScarlet + 1 mM ATP in the absence and presence of 5 nM sfGFP-MAP7 (means **±** s.d. are 0.09 **±** 0.11 vs. 1.34 **±** 0.43 motors per μm per min for K560 alone and K560 + MAP7, respectively; *P* < 0.0001; n = 28 events from 12 microtubules from two independent trials for K560 alone and n = 20 events from 8 microtubules from two independent trials for K560 + MAP7). All datapoints are plotted with lines indicating means **±** s.d. (**c**) Cumulative frequency distribution plot of K560-mScarlet run lengths (+ 1 mM ATP) in the absence and presence of sfGFP-MAP7 fit to a one phase exponential decay. Mean decay constants **±** s.d. are 873.8 **±** 43.3 and 984.3 **±** 87.5 nm for K560 and K560 + MAP7, respectively (*P* < 0.0001; n = 173 and 204 K560 motors for K560 alone and K560 + MAP7, respectively from three independent trials). (**d**) Velocity histograms of K560-mScarlet + 1 mM ATP in the absence and presence of sfGFP-MAP7 with Gaussian fits. Means **±** s.d. are 434.0 **±** 111.2 and 327.5 **±** 128.5 nm/sec for K560 alone and K560 + MAP7, respectively (*P* < 0.0001; n = 79 and 108 K560 motors for K560 alone and K560 + MAP7, respectively from three independent trials). (*e*) ATPase activities of 50 nM K560-mScarlet + 1 mM ATP in the absence and presence of 50 nM sfGFP-MAP7 as a function of microtubule concentration. All replicates are plotted from n = 3 independent experiments per condition and two separate protein preps and fitted with Michaelis-Menten kinetics (mean K_m_ **±** s.d. are 1.46 **±** 0.21 μM and 0.27 **±** 0.04 μM for K560 alone and K560 + MAP7, respectively; *P* = 0.0006). (**f**) Kymograph analysis of 10 nM K560-mScarlet + 1 mM ATP in the presence of 100 pM sfGFP-MAP7 reveals vertical, stationary MAP7 molecules (green) that do not co-migrate with processive K560 motors (pink diagonal lines). (**g**) TIRF-M images of 1 nM K560-mScarlet with 5 nM sfGFP wild type or mutant ensconsin recombinant proteins. Although mutant ensconsin proteins still bind the microtubule, they do not recruit K560 to the microtubule. Wild type ensconsin image is 10.3 μm in length and the mutant ensconsin images are 5.2 μm in length. (**h**) Quantification of the fold change in fluorescence intensity of K560-mScarlet on the microtubule in the presence of wild type, Δ665-699, or Δ693-699 ensconsin proteins. All datapoints are plotted with lines indicating means **±** s.d. (147.0 **±** 56.2, −6.5 **±** 20.9, and −2.5 **±** 13.4 fold intensity change for wild type, Δ665-699, or Δ693-699 ensconsin, respectively; n = 48, 53, and 51 microtubules, respectively from two independent trials). (**i**) TIRF-M images of different kinesin-1 constructs in the absence or presence of sfGFP-MAP7. Only K523 is recruited to the same extent as K560. All images are 9.5 μm in length. (**j**) Quantification of the fold change in fluorescence intensity of K490-, K508-, K523-, or K560-mScarlet on the microtubule in the presence of sfGFP-MAP7. All datapoints are plotted with lines indicating means **±** s.d. (−0.5 **±** 4.2, 10.9 **±** 4.0, 151.6 **±** 86.0, and 164.9 **±** 74.2 fold intensity change for K490, K508, K523, and K560, respectively; n = 20, 29, 48, and 63 microtubules, respectively from two independent trials). (**k**) TIRF-M images of 100 nM mTagBFP-tau (pink), 50 nM sfGFP-MAP7 and the max projection of 10 nM K560-mScarlet (orange) from a 5 min movie reveal that K560 is recruited by MAP7 to the microtubule, but excluded from tau patches. All images are 10.3 μm in length. (**l**) ATPase activities of 50 nM K560-mScarlet + 1 mM ATP + 2 μM microtubules in the absence and presence of 500 nM mTagBFP-tau and 500 nM sfGFP-MAP7 reveals that tau inhibits the microtubule-stimulated ATPase activity of K560, but the addition of MAP7 rescues the ATPase activity of K560. All replicates are plotted from n = three independent experiments per condition and lines representing the means **±** s.d. (24.7 **±** 4.0 s^−1^ (n = 9), 2.9 **±** 0.8 s^−1^ (n = 8), and 21.8 **±** 3.7 s^−1^ (n = 7) for K560 alone, K560 + tau, and K560 + tau + MAP7, respectively; K560 vs. K560 + tau: *P* < 0.0001; K560 + tau vs. K560 + tau + MAP7: *P* < 0.0001; K560 vs. K560 + tau + MAP7: *P* = 0.1596). All experiments in Figure 4 were repeated with at least two separate protein preparations for each protein.

How might MAP7 enhance kinesin-1 binding to microtubules? MAP7 could recruit kinesin-1 to the microtubule and remain stably associated with the motor as it moves, acting as a tether, or MAP7 could recruit kinesin-1 through a transient interaction, releasing the motor once it begins to step along the microtubule. Additionally, MAP7 could indirectly recruit the motor by altering the microtubule lattice to favor motor binding. Dual-color single molecule experiments revealed that less than 1% of K560 molecules co-migrated with a MAP7 molecule (3/328 K560 molecules; Fig. 4f) under our conditions. This result suggests a recruitment, rather than a tethering role for MAP7 in enhancing kinesin-1 microtubule association.

We next examined if kinesin-1 directly interacts with MAP7. Solution-based pull-down assays revealed that both MAP7 and ensconsin modestly interact with K560 (Supplemental Fig. 2b-c), consistent with interaction studies done in cell lysates^17^. We turned to *Drosophila* ensconsin to refine the previously reported kinesin-1 binding domain, which spans 158 amino acids and contains a predicted coiled-coil domain^17^. Similar to human MAP7, *Drosophila* ensconsin also strongly recruited K560 to the microtubule (Fig. 4g-h). However, there is relatively low amino acid homology between MAP7 and ensconsin within this region (41% identity). Based on sequence conservation, we deleted aa 665-699 within ensconsin (Supplemental Fig. 2d). Removal of these amino acids did not affect microtubule association, but strongly perturbed the ability of ensconsin to recruit K560 to the microtubule (Fig. 4g-h). Excising the most conserved coiled-coil heptad within this region (693-699, +4 charge) also strongly disrupted ensconsin’s recruitment of K560 to the microtubule (Fig. 4g-h). Using truncation mutants of kinesin-1, we found that MAP7 did not recruit K490, modestly recruited K508 (11-fold), and recruited K523 similarly to K560 to microtubules (152-fold vs. 165-fold, Fig. 4i-j, Supplemental Fig. 2e). This region (500-523) has −5 charge and is conserved among eukaryotes, but is not conserved in *A. nidulans*, which lacks MAP7 (Supplemental Fig. 2f). The lack of effect on K490 suggests that MAP7 does not allosterically change the microtubule lattice to favor kinesin binding, indicating MAP7 must directly interact with kinesin-1 to facilitate the interaction between the kinesin-1 and the microtubule. Overall, our results suggest that MAP7 recruits kinesin-1 to the microtubule through a transient, ionic interaction between coiled-coil domains, but does not remain stably bound to kinesin-1 as it translocates along the microtubule, consistent with a previously proposed hypothesis^16^.

Having established that MAP7 and tau compete for microtubule binding and have differential effects on kinesin-1 transport, we examined how the presence of MAP7 influenced the behavior of kinesin-1 upon encountering a tau patch. We observed a similar percentage of K560 motors that detached, paused, or passed tau patches in the absence compared with the presence of MAP7 (Supplemental Fig. 3a-c), but MAP7 still heavily recruited K560 to the microtubule outside of tau patches (Fig. 4k). In addition, an excess of tau inhibited the microtubule-stimulated ATPase activity of K560, but the presence of MAP7 restored K560 ATPase activity (24.7 **±** 4.0, 2.9 **±** 0.8, and 21.9 **±** 3.7 s^−1^ for K560 alone, K560 + tau, and K560 + tau + MAP7, respectively; Fig. 4l). This result further supports a primarily transient role for MAP7 in the recruitment of kinesin-1 to the lattice, as opposed to a mechanism whereby MAP7 tethers kinesin-1 to the microtubule through a stable interaction.

We next investigated if MAP7 affected two other classes of processive transport microtubule motors: the dendritic and axonal cargo motor, kinesin-3 ^34^, and the ubiquitous minus-end directed dynein-dynactin motor complex (DDB)^35^. Single molecule assays revealed that, in contrast to kinesin-1, MAP7 substantially reduced the overall amount of kinesin-3 (KIF1A) on the microtubule by 4-fold and the number of kinesin-3 landing events on the microtubule ~5-fold (from 1.50 **±** 0.74 to 0.38 **±** 0.31 motors per μM per min. Fig. 5a-c). Tau also strongly inhibited kinesin-3 landing rate (0.24 **±** 0.24 motors per μM per min.) and motility similarly to its effects on kinesin-1 (Fig. 5b-d, Supplemental Fig. 3c). Interestingly, MAP7 had little effect on the overall amount of DDB on the microtubule or the velocity of DDB complexes (606.4 **±** 178.0 and 555.2 **±** 186.5 nm/sec for DDB alone and DDB + MAP7, respectively; Fig. 5e-g). Together, these results provide a framework for how MAPs dictate access to the microtubule and subsequently determine the spatial distribution of microtubule motors within a crowded intracellular environment.

**Figure 5:**
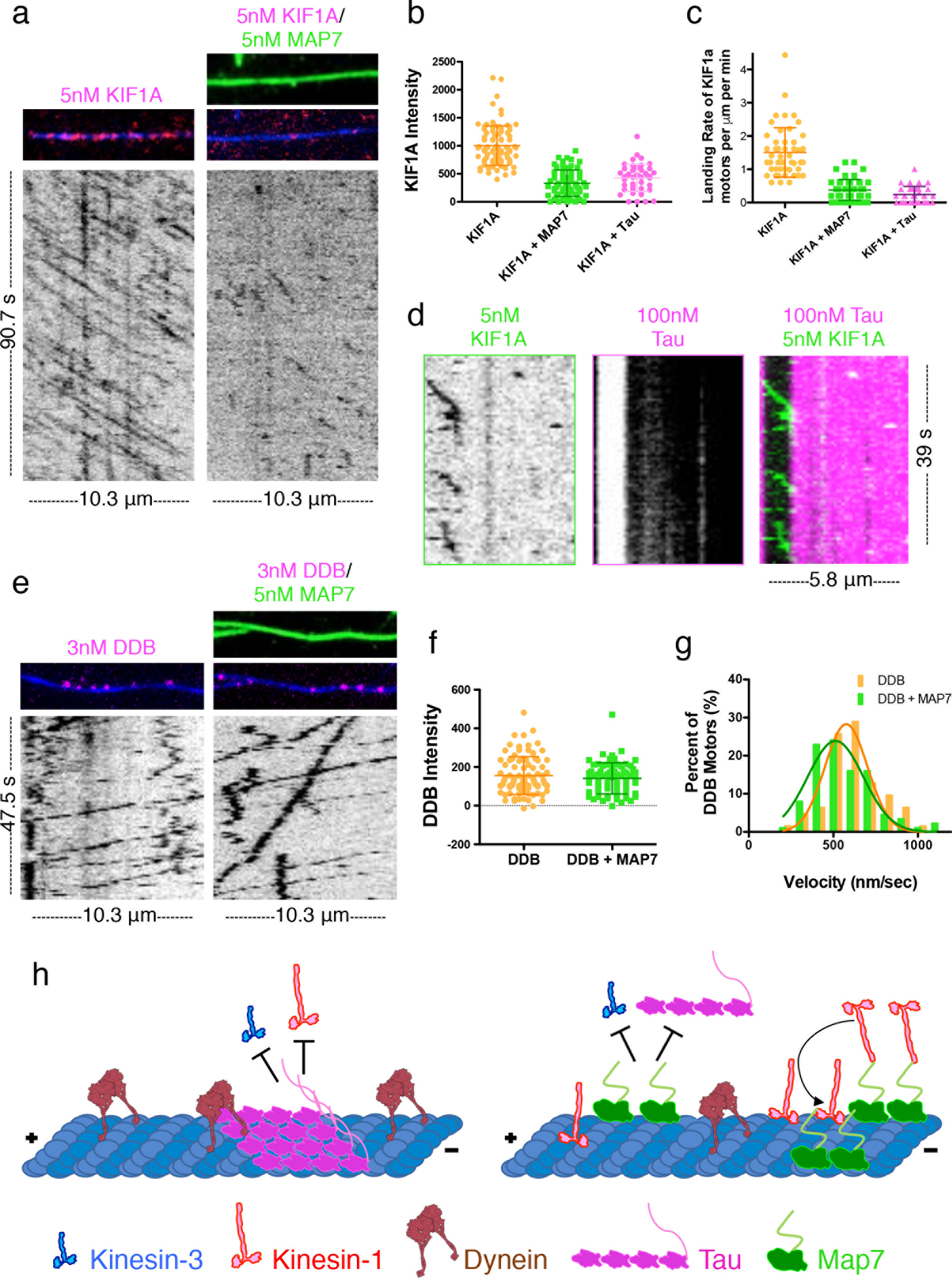
MAP7 and tau differentially affect other microtubule motors. (**a**) TIRF-M images and corresponding kymographs of 5 nM KIF1A-mScarlet (kinesin-3) in the absence and presence of 5 nM sfGFP-MAP7 reveal that MAP7 inhibits KIF1A from binding to the microtubule. Images are 10.3 μm in length. (**b**) Quantification of fluorescence intensity of KIF1A-mScarlet in the absence and presence of 5 nM sfGFP-MAP7 or 100 nM mTagBFP-tau (means **±** s.d. are 1003.1 **±** 351.1, 332.2 **±** 235.2, and 427.1 **±** 243.7 A.U., and n = 89, 69, and 46 microtubules for KIF1A alone, KIF1A + MAP7, and KIF1A + tau, respectively from three independent trials; *P* < 0.0001 for KIF1A vs. KIF1A + MAP7, as well as for KIF1A vs. KIF1A + tau). (**c**) Quantification of landing rates of KIF1A motors in the absence and presence of 5 nM sfGFP-MAP7 or 100 nM mTagBFP-tau (means **±** s.d. are 1.50 **±** 0.74, 0.38 **±** 0.31, and 0.24 **±** 0.24 motors per μm per min, and n = 50, 51, and 38 for KIF1A alone, KIF1A + MAP7, and KIF1A + tau, respectively from three independent trials and two separate protein preps; *P* < 0.0001 for KIF1A vs. KIF1A + MAP7, as well as for KIF1A vs. KIF1A + tau). Both graphs are scatterplots with all datapoints plotted and lines representing the means **±** s.d. (**d**) Kymograph depicting KIF1A-mScarlet motors encountering an mTagBFP-tau patch on the microtubule. (**e**) TIRF-M images and corresponding kymographs of 3 nM dynein-dynactin-BicD2 (DDB)-TMR + 1 mM ATP in the absence and presence of 5 nM sfGFP-MAP7. (**f**) Quantification of fluorescence intensity of 3 nM DDB-TMR in the absence and presence of 5 nM sfGFP-MAP7 (means **±** s.d. are 164.3 **±** 103.7 and 147.3 **±** 85.7 A.U. and n = 89 and 70 microtubules for DDB alone and DDB + MAP7, respectively from two independent trials; *P* = 0.27). (**g**) Velocity histograms of 3 nM DDB-TMR + 1 mM ATP in the absence and presence of 5 nM sfGFP-MAP7 with Gaussian fits. Means **±** s.d. are 606.4 **±** 178.0 and 555.2 **±** 186.5 nm/sec and n = 62 and 87 motors for DDB alone and DDB + MAP7, respectively from three independent trials (*P* = 0.095). All experiments in Figure 5 were repeated with at least two separate protein preparations for each protein. (**h**) Model for how competition between MAP7 and tau directs motor transport. In the presence of tau (pink), kinesin-1 (red) and kinesin-3 (blue) are inhibited from the microtubule, while dynein (burgundy) motility is unperturbed. The presence of MAP7 (green) evicts tau from the microtubule lattice, facilitating kinesin-1 recruitment and motility without altering dynein activity. MAP7 inhibits kinesin-3 similarly to tau, ensuring the exclusion of kinesin-3 from microtubules in tau- and MAP7-rich environments.

Overall, our study presents a detailed analysis of the competition between tau and MAP7 and a molecular dissection of the functional interaction between MAP7 and kinesin-1. Previous studies have shown that tau inhibits kinesin-1 without dramatically affecting dynein^6,7^, posing a transport problem within the axons of neuronal cells (Fig. 5h). We have found that MAP7 can facilitate kinesin-1 motility in two ways without significantly affecting dynein. First, MAP7 actively competes with and displaces tau from the microtubule, and second, MAP7 directly recruits kinesin-1 to the microtubule (Fig. 5h). We further show that both MAP7 and tau inhibit kinesin-3, which transports cargo into both dendrites and axons (Fig. 5h). The differential effects of these MAPs on microtubule motors not only establishes the balance of kinesin-1 and dynein transport in the axon, but may also help to direct kinesin-3 motors into the dendrites during specific developmental stages.

An imbalance of microtubule-based transport has been implicated in disease^9,10^, and our data suggest that competition between MAPs likely plays a role in coordinating the correct spatiotemporal activation of particular types of microtubule motors. In neurodegenerative diseases such as Alzheimer’s and fronto-temporal dementia, tau becomes hyperphosphorylated and dissociates from the microtubule^36^. In the absence of tau, the presence of MAP7 on the microtubule could tip the balance of transport in favor of kinesin-1, similar to what we observe upon overexpression of MAP7 in vivo (Fig. 3). This indirect disruption of dynein-based retrograde transport could have neurodegenerative effects, as has been seen in patients with mutations in dynein pathway components^9^.

Our results also raise intriguing questions about competition between other MAPs. The microtubule-binding domain of MAP7 is distinct from that of tau, MAP2, and MAP4, which contain similar tandem microtubule binding repeats^37^. Based on our results, it is possible that tau and MAP7 also compete with other MAPs for microtubule binding. Competition or coordination between MAP7 and dendrite-specific MAPs, such as MAP2 or DCX, may be especially important for directing kinesin-3 transport, because MAP7 is present in the dendrites, but inhibits kinesin-3 in vitro. Likewise, there may be axonal MAPs that enable kinesin-3 to drive transport in the axons in the presence of both MAP7 and tau. Additionally, there are other mechanisms of regulation that could dictate the spatiotemporal association patterns of MAPs on microtubules, such as post-translational modifications of tubulin^38^ or of the MAPs themselves. Overall, our results suggest that competition between MAPs may be a general mechanism for directing motor transport in cells. Combined with the tubulin-code hypothesis, which posits that modifications of the tubulin subunits themselves modulate molecular motor transport^38,39^, our data contributes another layer of regulation, the MAP-code, that we suggest may be similarly effectual in the spatiotemporal control of intracellular transport.

## ACKNOWLEDGEMENTS

We thank Ruensern Tan and Richard McKenney for providing the purified DDB complex, and Richard McKenney for useful discussion and critical reading of the manuscript. We thank Regis Giet for generously providing us with the ensconsin antibody and for useful discussion. We thank Scott Cameron and Kim McAllister for providing the fixed mouse neuronal cultures. The authors also thank the DRGC for providing the *ens* cDNA construct, and the Bloomington Drosophila Stock Center for providing fly stocks. This work was supported by the March of Dimes Basil O’Connor Award and NIH grant 1R00HD080981 to K.M.O.M. This material is based upon work supported by the National Science Foundation Graduate Research Fellowship Program under Grant No. 1650042 to B.Y.M. Any opinions, findings, and conclusions or recommendations expressed in this material are those of the authors and do not necessarily reflect the views of the National Science Foundation.

## AUTHOR CONTRIBUTIONS

B.Y.M. and K.M.O.M. designed the experiments and wrote the manuscript. B.Y.M., D.S., and T.T. purified the recombinant proteins. B.Y.M. performed the in vitro TIRF-M experiments. B.Y.M. and K.M.O.M. performed the neuronal culture, microtubule co-pelleting, and MAP7/Ens pull-down experiments and analyzed data. M.B. and K.M.O.M. performed the *Drosophila* work and analyzed data. D.S. and B.A. performed the ATPase assays and analyzed data.

---

**Supplementary Figure 1.**
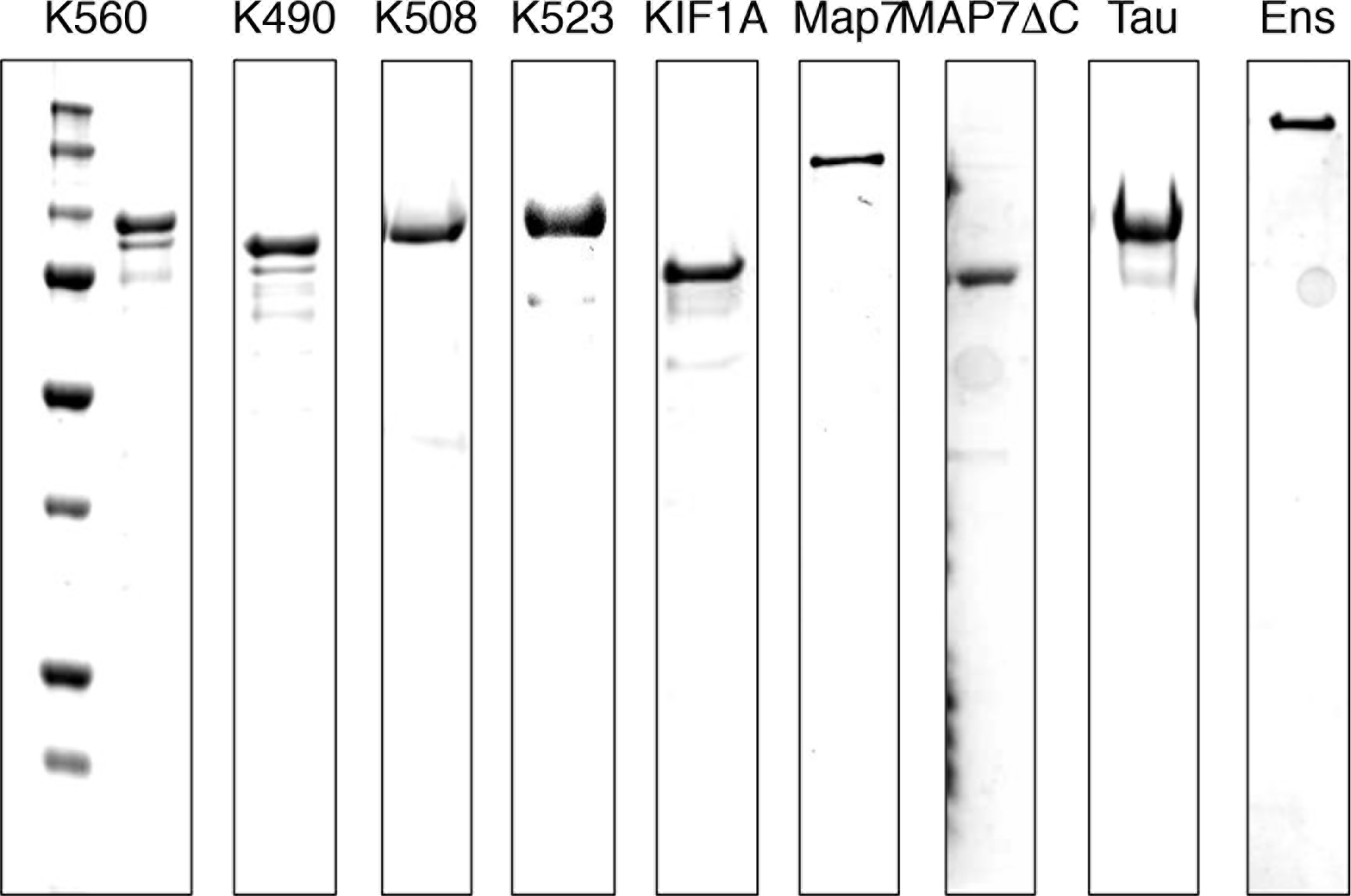
Purified recombinant proteins used in this study. Coomassie Blue-stained SDS-PAGE gels of K560-mScarlet, K490-mScarlet, K508-mScarlet, K523-mScarlet, KIF1A-mScarlet, sfGFP-MAP7, sfGFP-MAP7ΔC, mTagBFP-tau, and sfGFP-ensconsin.

**Supplementary Figure 2.**
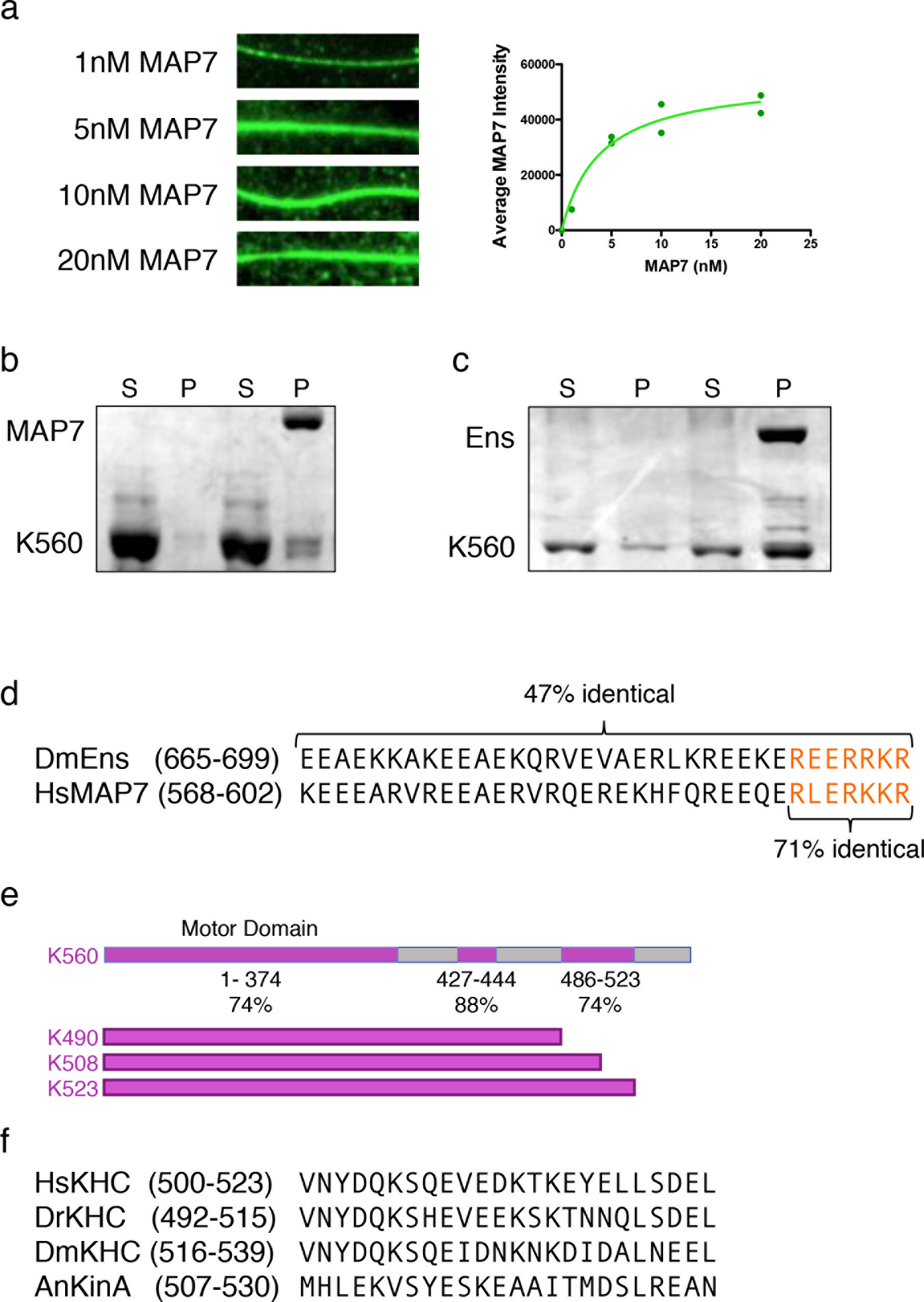
Dissection of the MAP7/ensconsin interaction with kinesin-1. (**a**) Fluorescence images of sfGFP-MAP7 bound to microtubules and corresponding quantification of fluorescence intensity of microtubule-bound sfGFP-MAP7 plotted against concentration (K_m_ = 3.8 **±** 1.2; n = 48, 21, 48, 44, and 49 microtubules for 0 nM, 1 nM, 5 nM, 10 nM, and 20 nM concentrations, respectively from two independent trials). (**b-c**) Coomassie Blue-stained SDS-PAGE gels of FLAG pull-downs with purified recombinant proteins. Either full-length sfGFP-MAP7-FLAG (**a**) or sfGFP-Ensconsin-FLAG were used for pull-down assays with K560-mScarlet (n = two independent trials per assay). S = supernatant and P = pellet. (**d**) Sequence alignment between *Drosophila melanogaster* ensconsin (DmEns) and *Homo sapiens* MAP7 (HsMAP7) with the percent identity indicated. (**e**) Schematic diagram of K560 and the associated constructs used in the mapping studies of Fig. 4i-j. The percent identities indicate the conservation between DmEns and HsMAP7. (**f**) Sequence alignment between HsKHC, *Danio rerio* KHC (DrKHC), DmKHC, and *Aspergillus nidulans* KinA (AnKinA).

**Supplementary Figure 3.**
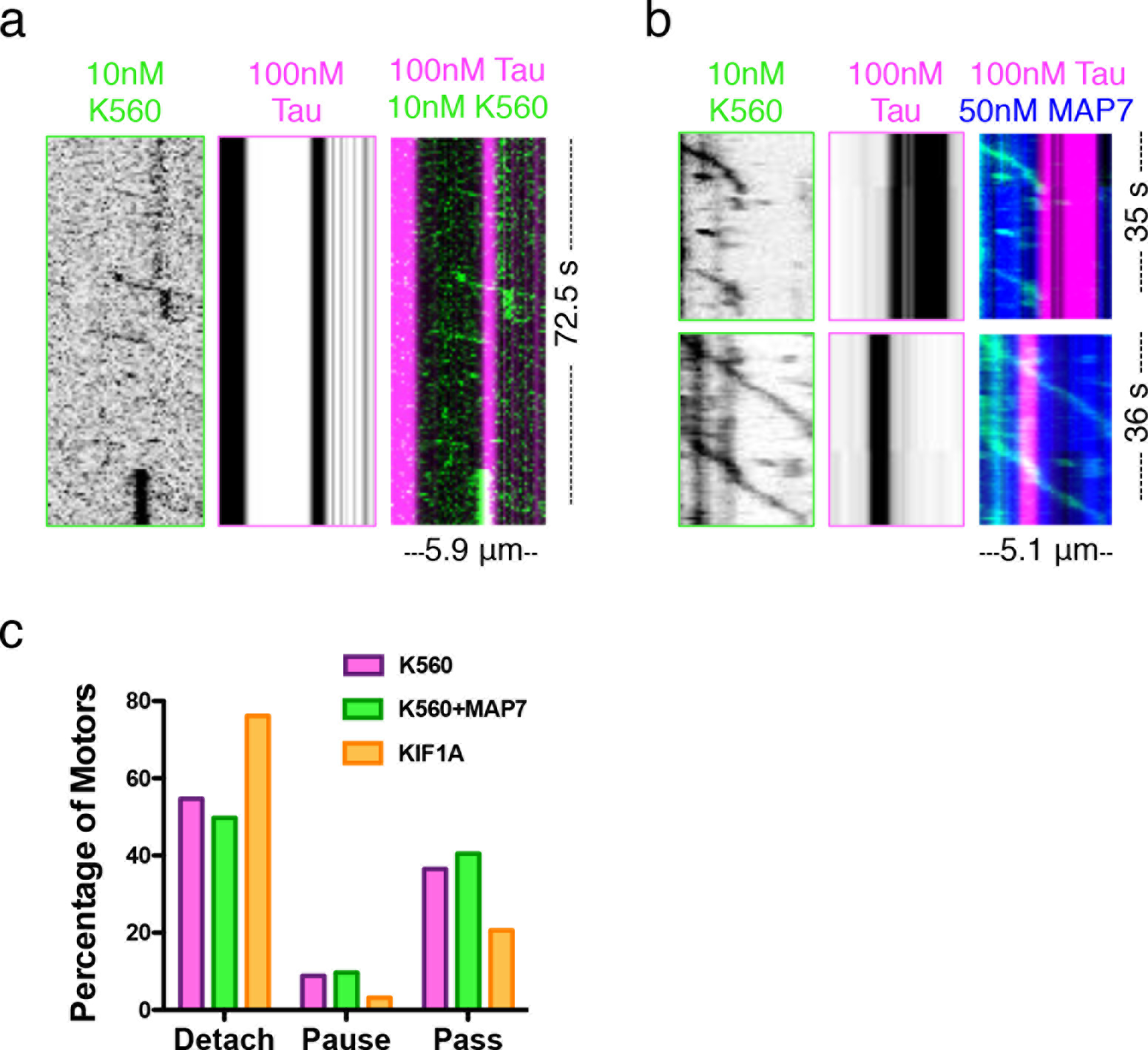
The effects of tau on kinesin-1 in the absence and presence of MAP7. (**a-b**) Kymographs depicting the effect of 100 nM mTagBFP-tau (pink) on 10 nM K560-mScarlet (green) motility in the absence (**a**) or presence (**b**) of 50 nM sfGFP-MAP7 (blue). (**c**) Quantification of the percent of K560 motors that detach, pause, or pass tau patches in the absence (pink) or presence (green) of MAP7 (54.7 % vs. 49.8 % detach, 8.8 % vs. 9.7 % pause, and 36.5 % vs. 40.5 % pass for K560 alone vs. K560 + MAP7, respectively; n = 159 from two independent experiments and 227 events from three independent experiments for K560 alone vs. K560 + MAP7, respectively). Related to Fig. 5d, also shown is the quantification of percent of KIF1A motors (orange) that detach, pause, or pass tau patches (76.2 % detach, 3.2 % pause, and 20.6 % pass; n = 126 motors from two independent experiments).

## METHODS

### Molecular Biology

The cDNAs for protein expression and fly injection used in this study were as follows: human Tau-2N4R (Addgene #16316), human MAP7 (GE Dharmacon MGC Collection #BC025777), human K560 (a gift from R. Vale), *Drosophila melanogaster* Ensconsin (Drosophila Genome Resource Center-DGRC #1070880), and human KIF1A (1-393) (Addgene # 61665). Tau-2N4R, MAP7, MAP7ΔC, and ensconsin proteins were cloned in frame using Gibson cloning into a pET28 or pFastBacHTA vector with an N-terminal strepII-Tag, mTagBFP, TagRFP, or a superfolder GFP (sfGFP) cassette. K560, K523, K508, K490, and KIF1A were cloned in frame using Gibson cloning into pET28 vector with a C-terminal mScarlet-strepII cassette.

### Protein Expression and Purification

Tubulin was isolated from porcine brain using the high-molarity PIPES procedure as previously described^40^. For bacterial expression of mTagBFP-Tau, sfGFP-Tau, sfGFP-MAP7, TagRFP-MAP7, sfGFP-MAP7AC, K560-mScarlet, K523-mScarlet, K508-mScarlet, K490-mScarlet, and KIF1A-mScarlet, BL21-RIPL cells were grown at 37°C until ~O.D. 0.6 and protein expression was induced with 0.1 mM IPTG. Cells were grown overnight at 18°C, harvested, and frozen. *Drosophila melanogaster* sfGFP-Ensconsin proteins were grown at 20°C for 4 hours before being harvested. Cell pellets were resuspended in lysis buffer (50 mM Tris pH 8, 150 mM K-acetate, 2 mM Mg-acetate, 1 mM EGTA, 10% glycerol) with protease inhibitor cocktail (Roche), 1 mM DTT, 1 mM PMSF, and DNAseI. Cells were then passed through an Emulsiflex press and cleared by centrifugation at 23,000 x *g* for 20 mins. For baculovirus expression of sfGFP-MAP7 or TagRFP-MAP7, the Bac-to-Bac protocol (Invitrogen) was followed. SF9 cells were grown in shaker flasks to ~2x10^6^/mL and infected at a ratio of 10mL virus to 250 mL cells. The infection was allowed to proceed for 48 hr before cells were harvested and frozen in LN_2_. Cell pellets were resuspended in lysis buffer (50mM Tris-HCl, pH 8.0, 150mM K-acetate, 2mM Mg-acetate, 1mM EDTA, 10% glycerol, 0.1mM ATP) with protease inhibitor cocktail (Roche). Cells were lysed by addition of 1% Triton X-100 for 10 min on ice. Clarified lysate from either bacterial or baculovirus expression was passed over a column with Streptactin Superflow resin (Qiagen). After incubation, the column was washed with four column volumes of lysis buffer, then bound proteins were eluted with 3 mM desthiobiotin (Sigma) in lysis buffer. Eluted proteins were concentrated on Amicon concentrators and passed through a superose-6 or superdex-200 (GE Healthcare) gel-filtration column in lysis buffer using a Bio-Rad NGC system. Peak fractions were collected, concentrated, and flash frozen in LN_2_. Protein concentration was determined by measuring the absorbance of the fluorescent protein tag and calculated using the molar extinction coefficient of the tag. Purified BicD2N was used to isolate DDB complexes from rat brain cytosol as previously described^35^. DDB complexes were labeled with 5 μm SNAP-TMR dye during the isolation procedure, and were frozen in small aliquots and stored at −80°C. The resulting preparations were analyzed by SDS-PAGE.

### Co-sedimentation Assays

Co-sedimentation assays were performed as previously described^41^. Microtubules were prepared by polymerizing 25 mg/mL of porcine tubulin in assembly buffer (BRB80 buffer supplemented with 1mM GTP, 1mM DTT) at 37°C for 15 min, then a final concentration of 20 μm taxol was added to the solution, which was incubated at 37°C for an additional 15 min. Microtubules were pelleted over a 25 % sucrose cushion at 100,000g at 25°C for 10 min, then resuspended resuspended in BRB80 buffer with 1mM DTT and 10 μm taxol. Binding reactions were performed by mixing 500 nM of sfGFP-MAP7 or sfGFP-tau (that had been pre-centrifuged at 100,000g) with the indicated concentrations of microtubules in assay buffer (50 mM Tris pH 8, 150 mM K-acetate, 2 mM Mg-acetate, 1 mM EGTA, 10% glycerol and supplemented with 1 mM DTT, 10 μm taxol, and 0.01 mg/mL BSA) and incubated at 25°C for 20 min. The mixtures were then pelleted at 90,000 x *g* at 25°C for 10 min. Supernatant and pellet fractions were recovered, resuspended in sample buffer, and analyzed by SDS-PAGE. Protein band intensities were quantified using ImageJ.

### Pull-down Assays

Pull-down assays were performed with either sfGFP-MAP7 or sfGFP-ensconsin tagged at the C-terminal end with a FLAG epitope. FLAG beads (Thermofisher) were washed into assay buffer (50 mM Tris, pH 8, 150 mM K-acetate, 2 mM Mg-acetate, 1 mM EGTA, 10% glycerol and supplemented with 1 mM DTT and 0.01 mg/mL BSA), then incubated with 500 nM MAP7, 500 nM ensconsin, or buffer (beads alone control) for 1 hour rotating at 4°C. FLAG beads were then washed in assay buffer five times, then resuspended in assay buffer and 500 nM (for ensconsin) or 1 μm K560 (for MAP7) was added to the beads alone control and the experimental condition. The 350 μL final volume solutions were incubated for 1 hour rotating at 4°C. The supernatants were collected, then the bead pellets were washed five times in assay buffer and resuspended in one bead bed volume. Gel samples of the supernatants and pellets were analyzed by SDS-PAGE. The supernatant samples that were run on SDS-PAGE were 15% of the pellet samples.

### TIRF Microscopy

TIRF-M experiments were performed as previously described^35^. A mixture of native tubulin, biotin-tubulin, and fluorescent-tubulin purified from porcine brain (~10:1:1 ratio) was assembled in BRB80 buffer (80mM PIPES, 1mM MgCh, 1mM EGTA, pH 6.8 with KOH) with 1mM GTP for 15 min at 37°C, then polymerized MTs were stabilized with 20 μm taxol. Microtubules were pelleted over a 25% sucrose cushion in BRB80 buffer to remove unpolymerized tubulin.

Flow chambers containing immobilized microtubules were assembled as described^35^. Imaging was performed on a Nikon Eclipse TE200-E microscope equipped with an Andor iXon EM CCD camera, a 100X, 1.49 NA objective, four laser lines (405, 491, 568, and 647 nm) and Micro-Manager software ^42^. All MAP7 and tau competition experiments were performed in assay buffer (30 mM Hepes pH 7.4, 150 mM K-acetate, 2 mM Mg-acetate, 1 mM EGTA, and 10% glycerol) supplemented with 0.1mg/mL biotin-BSA, 0.5% Pluronic F-168, and 0.2 mg/mL κ-casein (Sigma).

For all competition experiments, one protein was held at a constant concentration (50 nM sfGFP- or TagRFP-MAP7 or 100 nM mTagBFP- or sfGFP-tau) while the other was increased; both proteins were premixed and flowed into the chamber at the same time. Competition experiments were performed with a mixture of either sfGFP-MAP7 and mTagBFP-tau or a mixture of TagRFP-MAP7 and sfGFP-tau. For live imaging, images were taken every 15 seconds for a total of 10 minutes. For fluorescence intensity analysis, ImageJ was used to draw a line across the microtubule of either the Tau or MAP7 channel and the integrated density was measured. The line was then moved adjacent to the microtubule of interest and the local background was recorded. The background value was then subtracted from the value of interest to give a corrected intensity measurement. For dwell times, either MAP7 or tau was diluted to single molecule levels in the above-mentioned assay buffer and imaged at 0.5 s/frame and 0.128 sec/frame, respectively. The dwell times were measured by kymograph analysis in ImageJ by drawing a rectangle box around the beginning and end of a single dwelling molecule. The cumulative frequencies of MAP7 and tau dwell times were fit to a one phase exponential decay to derive the decay constants (*τ*) for each MAP.

All MAP7-kinesin construct motility and recruitment assays were performed in BRB80 buffer (80 mM PIPES pH 6.8, 1 mM MgCl_2_ and 1 mM EGTA) supplemented with 1 mM ATP, 150 mM K-acetate, with 0.1mg/mL biotin-BSA, 0.5% Pluronic F-168, and 0.2 mg/mL κ-casein. MAP7 was flowed in first, followed by K560, K523, K508 and K490. Fluorescence intensity analysis was performed as described above. Kymographs were made from movies of K560 in the absence and presence of MAP7 and velocity and processivity parameters were measured for individual K560 runs that moved processively for ≥ 250 nm at a speed of ≥ 80 nm/sec. Velocity data were fit with a Gaussian equation and the cumulative frequencies of K560 run lengths were fit to a one phase exponential decay to derive the mean decay constant (*τ*).

MAP7-tau-K560 experiments were carried out in assay buffer (30 mM Hepes pH 7.4,
150 mM K-acetate, 2 mM Mg-acetate, 1 mM EGTA, and 10% glycerol) supplemented with 1 mM ATP, 0.1mg/mL biotin-BSA, 0.5% Pluronic F-168, and 0.2 mg/mL κ-casein. MAP7 and tau were flowed in simultaneously, followed by K560. Kymographs were analyzed at regions of microtubules where there were clear tau patches that K560 molecules encountered. Events of pausing, passing, or detaching from the tau patch were counted. MAP7-KIF1A, tau-KIF1A, and MAP7-DDB experiments were performed in assay buffer (30 mM Hepes pH 7.4, 150 mM K-acetate, 2 mM Mg-acetate, 1 mM EGTA, and 10% glycerol) supplemented with 1 mM ATP, 0.1mg/mL biotin-BSA, 0.5% Pluronic F-168, and 0.2 mg/mL κ-casein. Fluorescence intensity analysis and velocity measurements were performed as described above. Events of pausing, passing, or detaching from the tau patch were counted for KIF1A.

### Fly Stocks

We used the Gal4 driver line, *221-Gal4^27^*, to drive expression of *UAS-CD4-RFP^27^* to visualize the dendrite morphology of class I DA neurons, UAS-*manII-eGFP^27^* to visualize Golgi outposts, UAS-*ensconsin-mCardinal* (this study) and UAS-*GFP-tau^43^*. We generated the UAS-*ensconsin-mCardinal* fly line by cloning the full-length ensconsin gene from the DGRC (#1070880) into a pUASt vector with an mCardinal tag at the C-terminus. Injection services were performed by Rainbow Transgenics. The endogenous Mi[MIC] GFP-tau line (Stock 60199) was obtained from the Bloomington Drosophila Stock Center (Department of Biology, Indiana University, Bloomington, IN).

### Live Imaging and Analysis

Whole, live third instar larvae were mounted in 90% glycerol under coverslips sealed with grease, and imaged using an Olympus FV1000 laser scanning confocal microscope. Live imaging and analysis were performed as previously described^25,28^. For morphological and Golgi outpost distribution analysis, the following genotypes were imaged: 1) *221-Gal4, UAS-CD4-RFP*, 2) *221-Gal4, UAS-CD4-RFP; UAS-GFP-tau*, 3) *221-Gal4, UAS-CD4-RFP; UAS-ens-mCardinal*, 4) *221-Gal4, UAS-CD4-RFP; UAS-GFP-tau/UAS-ens-mCardinal*, 5) *221-Gal4, UAS-CD4-RFP;UAS-manII-eGFP*, and 6) *221-Gal4, UAS-CD4-RFP;UAS-manII-eGFP/UAS-ens-mCardinal*. Z-stacks containing the dendritic arbors of the class I DA sensory neurons were collected for analysis from segment A3 or A4. Total dendrite length and number of branchpoints were determined from maximum Z-projections of the Z-stack image files using the Simple Neurite Tracer plugin ^44^ for ImageJ Fiji.

### Immunohistochemistry and Immunocytochemistry

Immunohistochemical staining on third larval instar fillets was performed as reported previously^25^. The primary antibodies used were chicken anti-GFP (GFP-1020, Aves Labs, RRID:AB_10000240), goat anti-HRP-Cy5 (123-605-021, Jackson ImmunoResearch, RRID:AB_2338967) and rabbit anti-ensconsin 172 (gift from R. Giet). For the ensconsin antibody staining, it was essential that larval muscle be cleared entirely from the fillet in order to get staining of the neurons, because this antibody prominently stains the muscles. Secondary antibodies consisted of appropriate fluorescence-conjugated anti-donkey IgG (Jackson ImmunoResearch). Slides were imaged on an Olympus FV1000 laser scanning confocal microscope using an oil immersion 40x or 60x objective.

Neuronal cultures were isolated from mouse cortices and cultured for 4 days in vitro (DIV) as previously described^41^. The cultures were fixed in 4% paraformaldehyde for 20 minutes at room temperature, washed several times with PBS, permeabilized in PBS with 0.3% Triton X-100 (PBS-TX), and blocked with 5% BSA in PBS for 1 hour at room temperature. Cultures were then incubated overnight at 4°C with primary antibodies at a concentration of 1:300 for rabbit anti-MAP7 (Thermofisher PA5-31782), 1:500 for mouse monoclonal anti-Tau (Millipore MAB3420), 1:1000 for mouse anti-alpha Tubulin (Sigma Clone DM1A T9026), or 1:1000 for rabbit anti-beta Tubulin (Abcam ab6046). Secondary antibodies were used at 1:1000 for Cy3 donkey anti-rabbit or Cy5 donkey anti-mouse and incubated for 1 hour at room temperature. Cells were then rinsed several times with PBS and mounted using VectaShield mounting medium (Vector Laboratories).

### ATPase Assays

ATPase assays were performed using an ATP/NADH coupled method in assay buffer containing 80 mM PIPES, pH 6.8, 1 mM MgCl2 and 1 mM EGTA, supplemented with 0.1% Triton-X, 1mM DTT, and 0.1 mg/mL BSA. The A340 was measured at 37°C for 5 minutes in the presence of assay buffer with 50nM K560, 2mM ATP, 0.2mM NADH (Roche), 2mM PEP (in 8mM KOH), 0.02% PK/LDH (enzymes from rabbit muscle, Sigma), and at a range of taxol-stabilized microtubule concentrations. Both mTagBFP-tau and sfGFP-tau were used in this assay, as well as sfGFP-MAP7 that was purified from insect cells using the baculovirus protocol. For the assays containing MAP7 and/or tau with K560, MAP7 and/or tau was incubated with microtubules for 5 minutes before K560 was added and the absorbance was measured. ATPase rate was measured and divided by the K560 (ATPase) concentration to determine the molar-activity of K560 per second. The ATPase rates were corrected for background NADH decomposition of controls containing no K560. MAP7 and tau did not exhibit ATPase activity on their own either in the presence of absence of microtubules. The resulting molar activity per second was plotted at a range of microtubule concentrations, and Michaelis-Menten curves were fit to the data to derive Km and Vmax values.

### Statistical Analysis

All statistical tests were performed with two-tailed Student’s t-test.

